# A cardiac pulse signal affects local field potentials recorded from deep brain stimulation electrodes across clinical targets

**DOI:** 10.64898/2026.02.10.704848

**Authors:** Katherine Tourigny, Rory J Piper, Martin Tisdall, Wolf-Julian Neumann, Alexander L Green, Timothy Denison, Joram van Rheede

**Affiliations:** Medical Research Council Centre of Research Excellence in Restorative Neural Dynamics, United Kingdom; Medical Research Council Brain Network Dynamics Unit, Nuffield Department of Clinical Neurosciences, University of Oxford, Level 6, West Wing, John Radcliffe Hospital, Oxford, United Kingdom, OX3 9DU; Division of Neurosurgery, Dalhousie University, Halifax Infirmary, 1796 Summer Street Halifax, NS, Canada, B3H 3A7; Developmental Neurosciences, UCL Great Ormond Street Institute of Child Health, University College London, 30 Guilford Street, London, United Kingdom, WC1N 1EH; Great Ormond Street Hospital for Children NHS Trust, Guilford Street, London, United Kingdom, WC1N 3BH; Movement Disorder and Neuromodulation Unit, Department of Neurology, Charité – Universitätsmedizin Berlin, corporate member of Freie Universität Berlin and Humboldt-Universität zu Berlin, Chariteplatz 1, 10117 Berlin, Berlin, Germany; Nuffield Department of Surgical Sciences, University of Oxford, John Radcliffe Hospital, Headington, Oxford, OX3 9DU; Institute of Biomedical Engineering, Department of Engineering Science, University of Oxford, Old Road Campus Research Building, Headington, Oxford OX3 7DQ

**Keywords:** Local field potential, deep brain stimulation, artifacts, neuromodulation, signal processing

## Abstract

**Objective:** Many deep brain stimulation (DBS) systems sense local field potentials (LFPs) for patient monitoring or closed-loop therapy (CL-DBS). LFPs can be impacted by artifacts, including a recently discovered cardiac non-electrocardiographic pulse signal that can be visually masked by commercial device filters. We aimed to establish its prevalence across patient groups and brain areas, and to investigate its spectral impact.

**Methods:** We performed a cross-sectional analysis of LFPs recorded from the cranially mounted Picostim from the pedunculopontine nucleus in multiple systems atrophy patients, periacqueductal gray and sensory thalamus in chronic pain patients, and the centromedian thalamic nucleus (CMT) in paediatric epilepsy patients. For comparison, we analyse externalised recordings from the subthalamic nucleus in Parkinson’s disease patients. The PulsAr algorithm was developed to detect and extract pulsatile signals, and we characterised contamination level and spectral content.

**Results:** Though not visually obvious in CMT, the pulsatile signal was algorithmically detected in all targets, with 33% of LFPs across targets classed as contaminated. Pulse signal power was similar across targets and may have been masked by higher endogenous activity in CMT. While its dominant frequencies were in the heartbeat range, the signal had spectral content extending up to >10Hz.

**Conclusions:** A heart pulse signal affects LFP recordings from DBS leads across brain regions and patient groups. While masked by some device filters, spectral content can extend into higher (clinically relevant) frequencies. Researchers and clinicians should exercise caution when sensing lower LFP frequencies, especially for automated control of therapy in CL-DBS.

**Highlights:**

- Historically, electrocardiographic artifacts have been a major source of artifact affecting deep brain stimulation recordings, however pulsatile artifact is less well described.
- A cardiac pulse signal affects local field potentials recorded from deep brain stimulation electrodes across clinical targets.
- We introduce an ECG-independent algorithm that detects and extracts this pulsatile signal.
- The heart pulse signal looks like the intracranial pressure waveform and affects spectral frequencies above the heart rate range up to >10Hz.
- Clinicians should incorporate screening and procedures to ensure accurate biomarker detection for clinical decision making.

## INTRODUCTION

Deep brain stimulation (DBS) improves symptoms of neurological disorders by modulating pathological neural circuits. It is an established therapy for movement disorders such as Parkinson’s disease (PD), dystonia, and essential tremor and for medically refractory epilepsy, with active research underway for other indications [1]. Many DBS devices now incorporate sensing technology to track neural activity around the therapeutic target. In closed loop DBS (CL-DBS), such activity is used to automatically adjust stimulation based on detected neural biomarkers, with the potential to improve efficacy and reduce side effects by maintaining neural activity within an optimal range [1,2]. CL-DBS or adaptive DBS (aDBS) systems have now been approved for human clinical use in epilepsy (NeuroPace Responsive Neurostimulation) and movement disorders (BrainSense adaptive DBS on the Medtronic Percept) [3–8]. In these systems, the accuracy of the “sensed” neural signal is crucial for achieving effective and safe control of stimulation [9].

Sensing-enabled DBS systems measure local field potentials (LFPs), representing the summed electrical activity of a population of neurons [10]. Oscillatory neural activity is often characterized using ‘canonical’ frequency bands of interest (FOIs) including slow oscillations (< 1 Hz), delta (1–4 Hz), theta (> 4–8 Hz), alpha (> 8–13 Hz), beta (> 13–30 Hz), and gamma (30–80 Hz)[11]. Certain oscillations have become established neurophysiological biomarkers, such as excessive beta activity in the subthalamic nucleus (STN) in PD which correlates with motor symptoms [12]. Ideally, LFPs are recorded, biomarkers are extracted, and stimulation is adjusted accordingly. However, artifact in LFPs can disrupt biomarker detection and feedback control (figure 1) [13].

**Figure 1.**
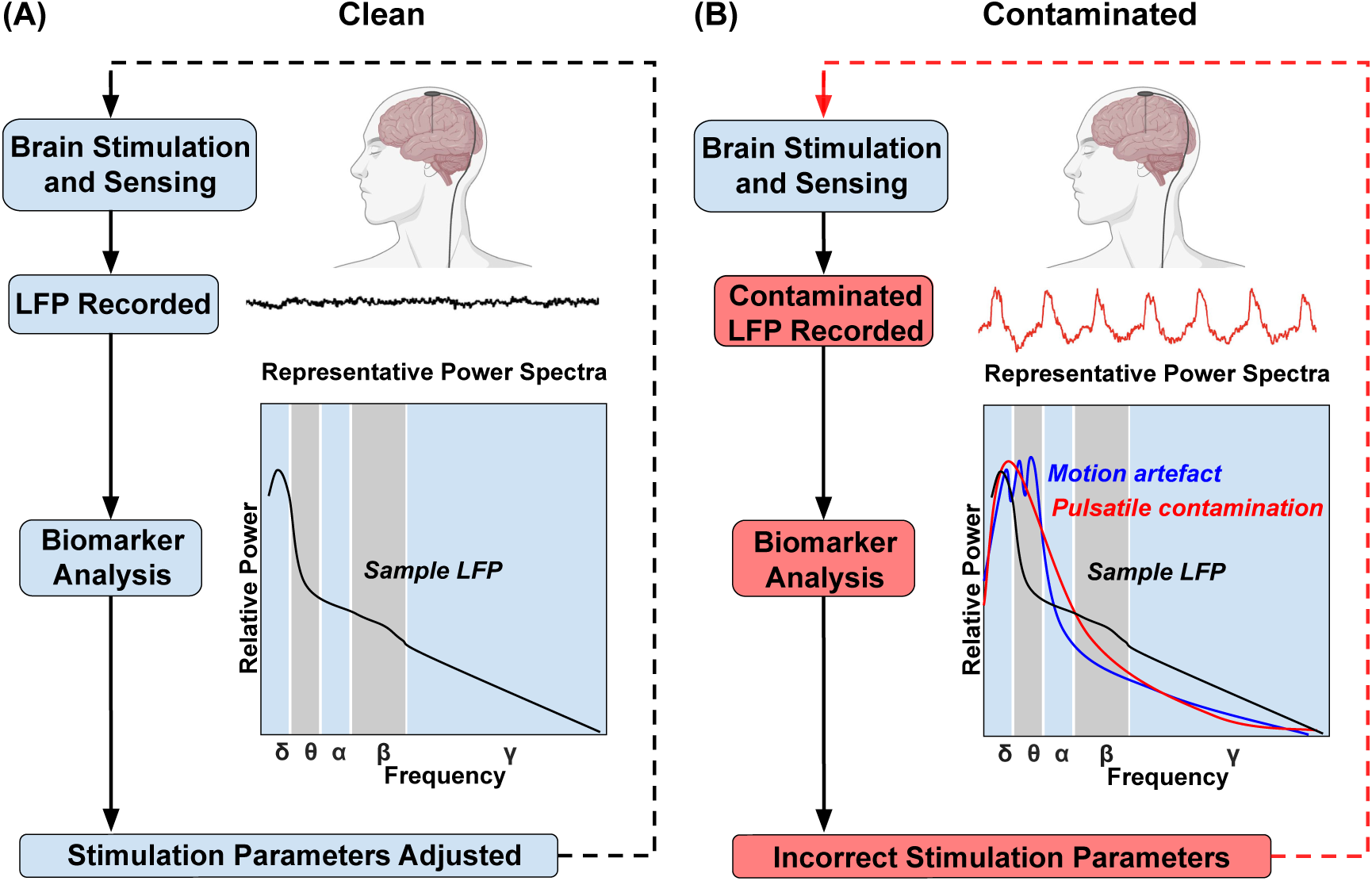
Closed loop DBS with clean and contaminated LFPs. (A) A clean LFP is recorded, and frequency bands of interest are analysed for biomarkers. Biomarker levels inform changes to the stimulation parameters through onboard biomarker analysis. New stimulation parameters are sent back to the brain. (B) The LFP is recorded and contaminated by artifact. An example of pulsatile signal is shown in the LFP. Schematic power spectra of a sample LFP, motion artifact, and pulsatile contamination. The power spectra of the artifact may overlap with frequency bands of interest. The onboard analysis may not be able to modify stimulation parameters, may revert to basal stimulation parameters, or may shut down as a safety mechanism.

Artifacts in neural signals originate from non-physiologic (e.g. environment, hardware, stimulation) or physiologic sources, and cardiac electrical activity is a well-known source of physiological artifact[14–19]. Electrocardiographic (ECG) contamination with the characteristic ‘QRS’ shape of the heart’s electrical field is well described, and several methods have been proposed for its removal [16–19]. QRS artifact can arise when the IPG is implanted close to the cardiac dipole and the QRS signal is not rejected as a common model [13]. Modelling studies predicted that ECG QRS contamination should not affect a cranially mounted IPG such as the Picostim™-DyNeuMo (“Picostim”) [13,17,20,21]. However, LFPs recorded with the Picostim in the pedunculopontine nucleus showed a large amplitude repetitive pulse-like signal temporally aligned with ECG but distinct from the well described QRS morphology [22]. Similar repetitive signals have been observed in intraoperative recordings [19]. Leading commercial devices can visually mask this pulse signal in LFP streams, with default high-pass filter cutoffs ranging from e.g. 1-10 Hz (Medtronic Percept, configurable) or 4-12 Hz (Neuropace RNS) [23–25]. However, this might not remove higher frequency components associated with dynamics of the cardiac cycle, which could mask neural signals relevant to therapy. The prevalence of this heart pulse signal in LFP recordings across the brain remains unclear.

In this paper, we characterize this pulsatile signal in multiple DBS targets and clinical populations and introduce an algorithm to detect it without concurrent ECG measurement. Next, we estimate its spectral content to predict its impact on biomarker frequency bands relevant to (CL-)DBS and assess its stability over time.

## METHODS

### Data Collection

We performed a retrospective analysis of prospectively collected local field potentials (LFPs) from participants enrolled in adult clinical trials of deep brain stimulation (DBS) of the pedunculopontine nucleus (PPN) for multiple system atrophy (n = 3, NCT05197816), periaqueductal gray (PAG) and sensory thalamus (ST) DBS for chronic post-stroke pain (n = 3, NCT06387914), as well as a pediatric trial of centromedian nucleus of the thalamus (CMT) DBS for Lennox–Gastaut epilepsy syndrome (n = 3, NCT05437393). These trials utilised the Picostim™-DyNeuMo (referred to as “Picostim”) cranially mounted adaptive DBS system (7.4-cc titanium IPG; two 4-contact leads). All LFPs available at the time of analysis were included.

Additional subthalamic nucleus (STN) recordings from one adult PD participant were obtained from externalized Medtronic 3389 or SenSight electrodes (Model B33005). Data collection followed each trial’s protocol as approved by local ethics boards and regulatory authorities and complied with the Declaration of Helsinki.

LFPs from the PPN, PAG, ST, and CMT were collected using a systematic montage sweep protocol (figure 2). Each sweep consisted of twenty-second bipolar recordings for all possible “within lead” contact pairs with one contact the anode and the other the cathode (contacts 1-4, or 5-8). Longer continuous recordings were available to assess temporal changes in the signal. Picostim recordings were sampled at 1000 Hz. STN recordings were collected from externalized leads at a sampling frequency of 4000 Hz using a Saga64 + (Twente Medical Systems International) device. STN signals were hardware-referenced to the lowermost contact of the DBS electrode and subsequently re-referenced offline in a bipolar configuration.

**Figure 2.**
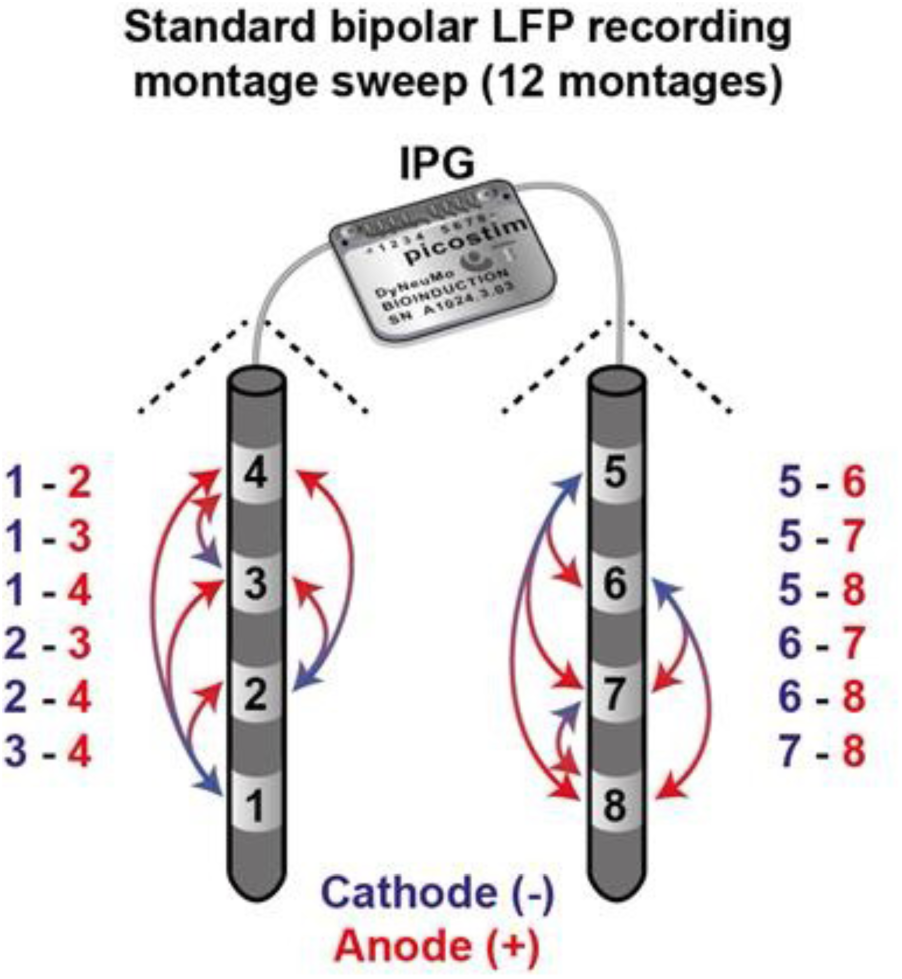
Schematic of the montage trace protocol using the Picostim system and a four-contact electrode. A twenty second LFP is recorded from all pairwise combinations of contacts sequentially in time. A standard bipolar sensing montage sweep consists of one contact serving as the anode and another as the cathode within the same electrode.

### Signal Processing

Analysis was performed in MATLAB (2023b, Mathworks Inc., Natick, MA, USA). LFPs were filtered with zero-phase, 4^th^ order Butterworth filters: 0.5 Hz high-pass and then a 100Hz low-pass filter. STN signals were then downsampled to 1000Hz. Power spectral densities (PSDs) for LFPs were computed using Welch’s method (MATLAB ‘*pwelch’*, 2000 sample windows, 50% overlap, 0.5 Hz resolution). Fast Fourier transforms (FFT) were used for pulsatile signal templates. A normalized spectral difference was computed at each frequency as the logarithmic ratio between contaminated and clean conditions. Wavelet spectrograms were computed with continuous Morlet wavelet transform (MATLAB *‘cwt’*) to allow for scaling of window length to more accurately capture changing frequency content within individual cardiac pulse cycles over time. Two raters (KT, JvR) visually classified traces as “clean” or “contaminated” based on the presence of a large amplitude repetitive pattern repeating at physiologic heart rate [16]. Signals were confirmed free of ECG QRS artifact using an open-source ECG detection and removal toolbox (Perceive, https://github.com/neuromodulation/perceive) [17]. We applied sequential high pass filters at 0.5 Hz, 1 Hz, 2 Hz, 4 Hz, 6 Hz, 8 Hz, 10 Hz, and 12 Hz to a pulsatile template string from a sample pulsatile template string from the PPN, and to the original contaminated PPN signal to assess high pass filter effect on visual assessment of signal contamination.

### PulsAr algorithm and pulsatile signal characteristics

We developed the PulsAr detection algorithm (“PulsAr”) to automatically classify LFPs as clean or contaminated. PulsAr uses cross-correlation to identify highly repetitive features in the signal. Raw LFPs were band-pass filtered between 0.5 Hz and 30 Hz using MATLAB’s *highpass* and *lowpass* filters to avoid phase distortion. Autocorrelation was computed for lags up to 10s. After discarding the first one second of signal to remove the zero-lag peak, the root-mean-square (RMS) power of the autocorrelation signal was computed as metric of contamination. Signals were classified as contaminated if they met three criteria: autocorrelation RMS power exceeded the threshold (see below explanation), inter-peak intervals were consistent with physiologic heart rates (50-120 bpm), and there was low variability in this inter-peak interval.

If the LFP was labelled as contaminated, a representative pulsatile template was extracted using a cross-correlation approach adapted from the Perceive toolbox [17]. An initial template was obtained from the autocorrelation peak, and this template cross-correlated with the original signal to identify segments sharing similar morphology. Peaks above 10% of the maximum correlation value and their locations were identified, segments around these peaks extracted, and segments averaged to generate a refined pulsatile template. The template was trimmed to zero-crossings and standardized to allow us to string together a repetitive “pulsatile artifact only” signal. This template was considered the representative pulsatile signal for each contaminated recording. Root mean square (RMS) amplitude was then compared across clean signals, contaminated signals, and pulsatile signal templates across brain regions.

PulsAr performance compared to experimenter ratings was evaluated using receiver operating characteristic (ROC) analysis and a confusion matrix, with consensus visual ratings from KT an JvR as ground truth. We ran PulsAr at 50 thresholds. The optimal threshold of 0.081 when compared to ground truth was used to classify signals for further analysis.

To assess stability of pulsatile signal over time, two PPN recordings over 40 seconds long were segmented into sliding 5-second windows. A pulsatile template was extracted by PulsAr for each 5-second window, and RMS amplitude calculated for each template to qualitatively evaluate within-recording variability.

### Statistical Analysis

Inter-rater agreement for visual classification was assessed using Cohen’s κ. Power spectral density (PSD) of clean and contaminated local field potentials (LFPs) was computed in linear units; group-level mean spectra and 95% confidence intervals (CI) were converted to decibels (dB) for visualization. Normalized spectral differences were calculated as the logarithmic ratio of contaminated to clean power. Band powers are reported as linear mean ± 95% CI. Frequency spectra of pulsatile templates are presented as the mean fast Fourier transform (FFT) ± 95% CI for each anatomical area. Differences between clean and contaminated signals were assessed using independent-samples t-tests with Welch’s correction, performed separately for each frequency band (delta, theta, alpha, beta, gamma), and corrected for multiple comparisons using the Benjamini–Hochberg (B-H) false discovery rate (FDR, q = 0.05). Root-mean-square (RMS) amplitude is reported as mean ± 95% CI for clean, contaminated, and pulsatile-template signals.

Group differences between clean and contaminated RMS amplitudes were assessed using Mann–Whitney U tests. Differences between pulsatile templates in each DBS target were assessed using the Kruskal–Wallis test. We calculated a contamination index to quantify the relative magnitude of pulsatile signal within contaminated LFP segments. The contamination index was defined as RMS of the pulsatile template over RMS of the signal and differences between groups assessed by Kruskal-Wallis test. The contamination index indicates the proportion of the total contaminated LFP signal amplitude attributable to pulsatile signal energy, with higher indices showing higher proportion of signal attributable to contamination. Boxplots show the median, 25th–75th percentiles (box), and minimum/maximum values within 1.5× inter-quartile range (whiskers). Outliers are shown as individual points.

## RESULTS

### Presence of a pulsatile artifact in intracranial LFP data

We visually inspected a total of 433 20 second montage sweep LFP recordings from the cranially mounted Picostim™-DyNeuMo for the presence of a pulsatile artifact. This included recordings from the pedunculopontine nucleus (PPN, n=154, MINDS trial), periacqueductal gray (PAG, n = 12, EPIONE trial), sensory thalamus (ST, n= 12, EPIONE trial), and centromedian thalamus (CMT, n = 255, CADET Pilot). Additionally, we investigated 30 subthalamic nucleus (STN) recordings obtained with externalized leads. Representative clean and contaminated signals from the PPN, PAG, ST, and STN, along with wavelet spectrograms of the contaminated examples, are shown in figure 3. Qualitative inspection showed a clear pulsatile signal in a proportion of PPN, PAG, ST, and STN recordings. In the PPN and PAG, spectrograms showed a prominent power at frequencies corresponding to physiologic heart rate, with spectral components extending into the 5-10Hz range. The ST and STN had similar but less pronounced patterns. In the PPN and PAG, a second smaller peak in the pulsatile signal can be appreciated. No clear pulsatile signal was seen in the CMT, although three traces were considered potentially contaminated by one rater.

**Figure 3.**
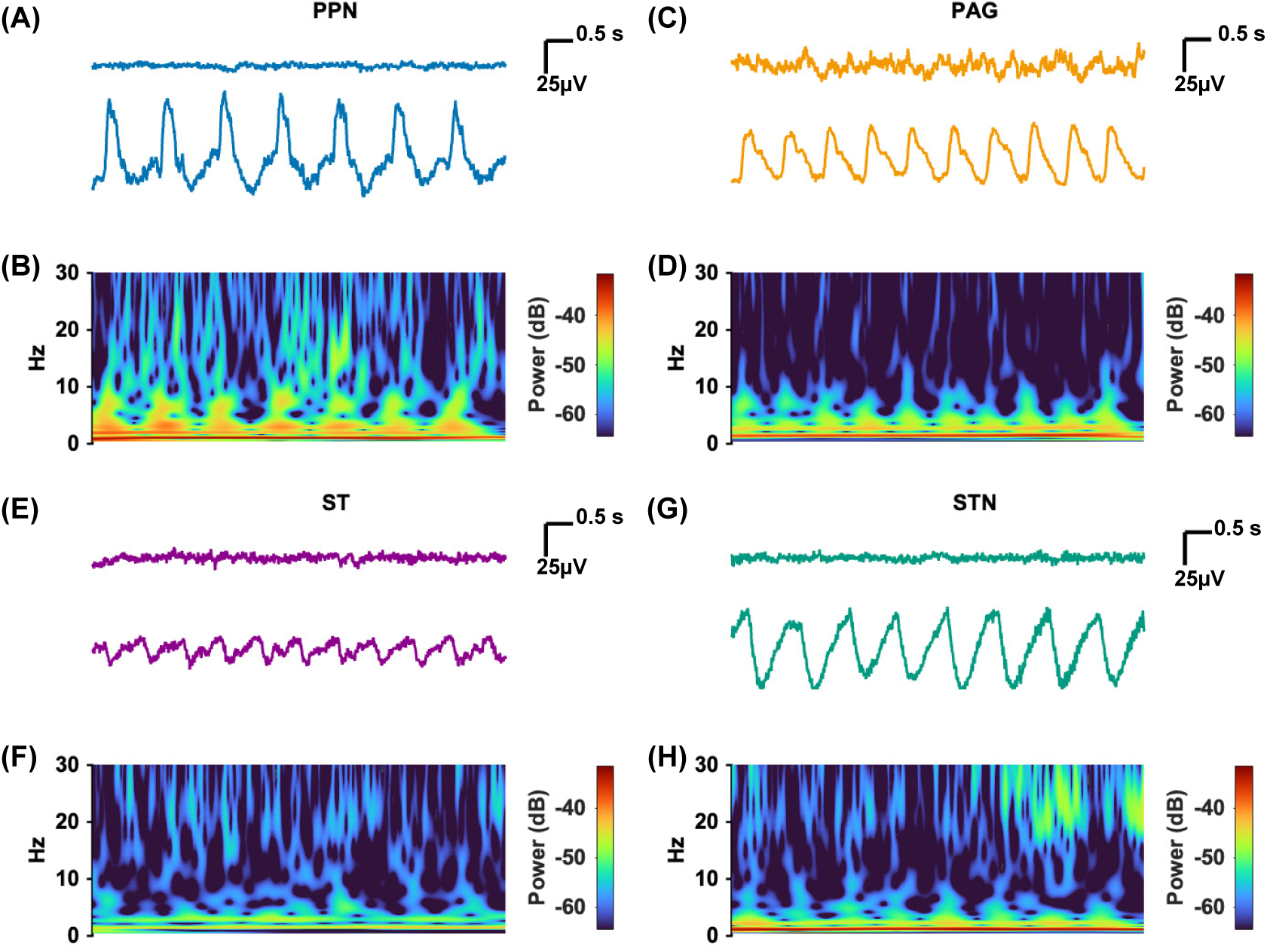
Sample clean and contaminated signals with wavelet spectrograms. Pulsatile signal morphology was similar across anatomic targets. (A) PPN clean (top) and contaminated (bottom) signals. (B) Wavelet spectrogram showing power of pulsatile signal (dB) in low frequencies and peaks temporally aligned with the peaks seen in the PPN LFP. (C)PAG clean (top) and contaminated (bottom) signals. (D) Wavelet spectrogram showing power of pulsatile signal (dB) in low frequencies and peaks temporally aligned with the peaks seen in the PAG LFP. (E) ST clean (top) and contaminated (bottom) signals. (F) Wavelet spectrogram showing power of pulsatile signal (dB) corresponding with the occurrence of pulsatile waveform in the LFP however less apparent compared to PAG and PPN. (G) STN clean (top) and contaminated (bottom) signals. (H) Wavelet spectrogram showing power of pulsatile signal (dB) corresponding with the occurrence of pulsatile waveform in the LFP however also less apparent than the PPN and PAG. Scalebars apply to all LFP and spectrogram panels. PPN = pedunculopontine nucleus, PAG = periaqueductal grey, ST = sensory thalamus, STN = subthalamic nucleus.

Across all participants, contamination did not necessarily affect all contact pairs within a montage sweep, and the same pair was not necessarily contaminated across different recording sessions. Overall, rater KT graded 69 traces as contaminated and JvR rated 114 as contaminated. Despite differences in rater conservativeness, inter-rater reliability was high (Cohen’s κ = 0.8).

We developed the PulsAr algorithm to enable objective detection and estimation of pulsatile artifact contamination (figure 4A-D; See Methods). As a ground truth for evaluating algorithm performance, a signal was labelled ‘contaminated’ if both raters considered a signal contaminated, and ‘clean’ otherwise. After evaluating 50 thresholds, a RMS contamination threshold of 0.081 optimized algorithm performance (sensitivity 94.2%, specificity 95.1%, F1 score 0.73, and AUC 0.92; figure 4E). This threshold was used to process the final refined dataset.

**Figure 4.**
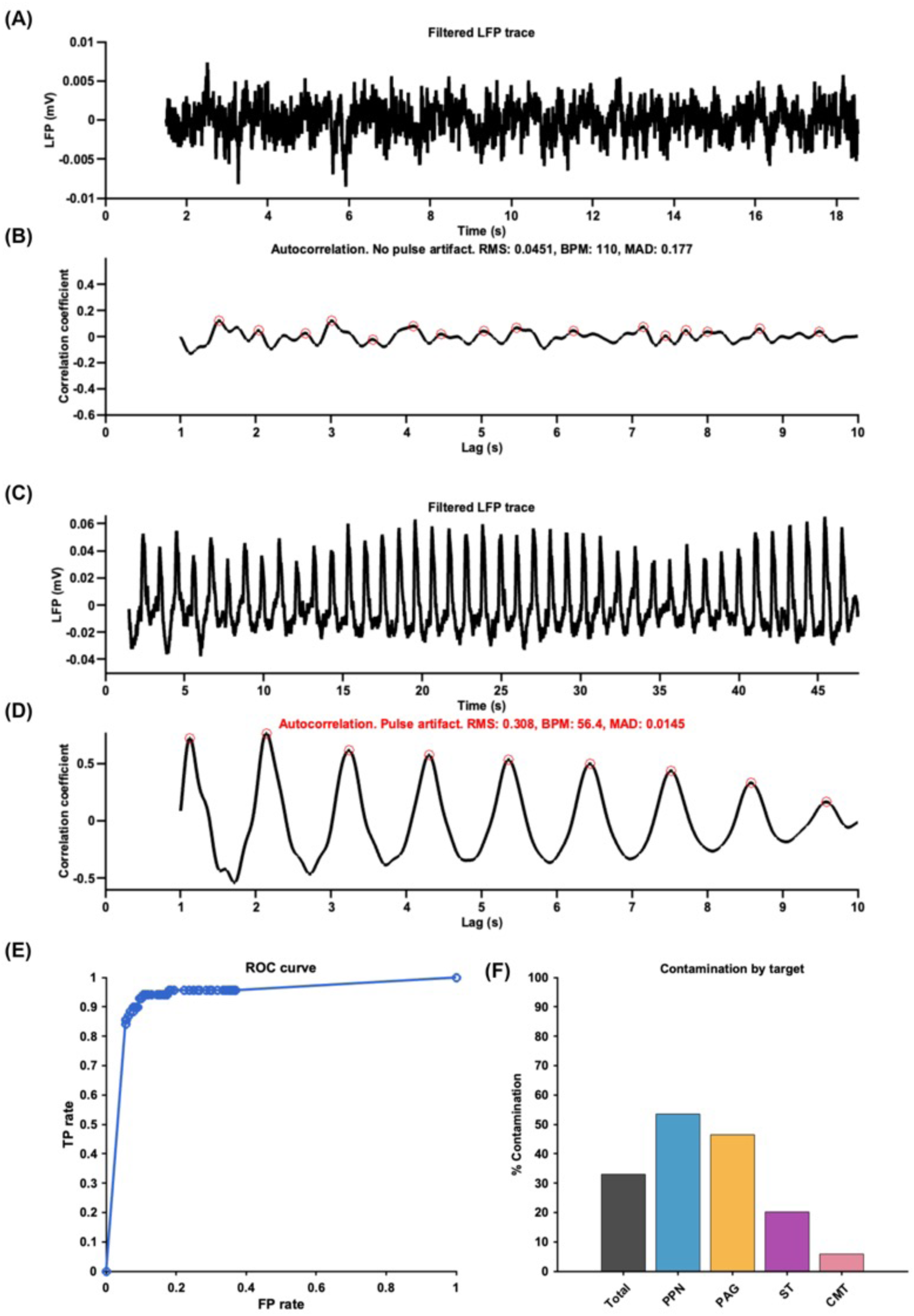
PulsAr output, signal contamination, and algorithm performance. (A-D) PulsAr algorithm outputs for a clean (A-B) and contaminated (C-D) signals. (A) Clean LFP from the sensory thalamus. (B) Autocorrelation showing no pulsatile signal detected. Autocorrelation step calculates root mean square power (RMS), estimated heart rate in beats per minute (BPM), and mean absolute deviation (MAD). (C) Contaminated LFP from the PPN showing large pulsatile signal. (D) Autocorrelation detected pulsatile signal. (E) Receiver operating characteristics (ROC) curve was calculated by comparing the PulsAr decision with consensus visual inspection from two raters KT and JvR. If the raters did not agree the consensus decision was to label the trace non-contaminated. The PulsAr algorithm was run at 50 different thresholds on all traces to build the ROC curve. (F) Percent of contaminated traces from all traces at each specific target. Overall, 33.0% of the LFPs collected from the Picostim were labelled as contaminated by PulsAr. Of all PPN LFPs 53.5% were contaminated, 44.0% of PAG LFPs, 22.6 % of ST LFPs, and 5.8% of CMT LFPs. RMS = root mean square power. BPM = beats per minute (estimated heart rate). MAD = mean absolute deviation.

The final dataset from the clinical trials included 445 montage sweep LFPs: 157 PPN, 84 PAG, 84 ST, and 120 CMT. LFPs from the STN (n = 30) were excluded from prevalence calculations as they were not collected under our standardized montage sweep parameters. The PulsAr algorithm detected contamination in 33% of all LFP recordings, including in 5.8% of CMT recordings which visually looked non-contaminated (Table 1). Sample PulsAr outputs for clean and contaminated signals, ROC, and % contamination results are shown in figure 4.

**TABLE 1.**
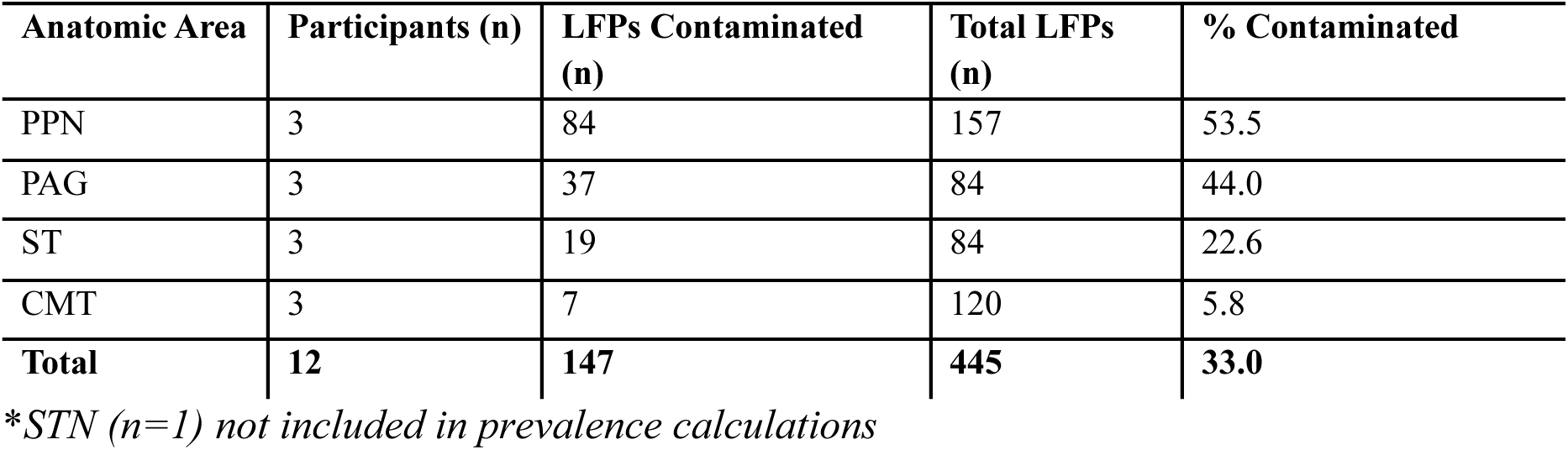
PulsAr Contamination Results.

We next examined how clean and contaminated signals differed in their spectral content. Contaminated signals displayed consistent increases in the lower frequency bands (figure 5). Comparison of power bands between clean versus contaminated signals (Table 2) revealed significant differences delta power in the PPN (B-H adjusted p = 0.021) and PAG (B-H adjusted p= 0.013). In the ST there were significant differences in beta (B-H adjusted p = 0.011) and gamma (B-H adjusted p = 0.011). In the CMT there was significantly lower spectral power in the theta (B-H adjusted p = 0.026) and alpha power (B-H adjusted p = 0.026) in contaminated traces. There were no significant differences in the STN.

**Figure 5.**
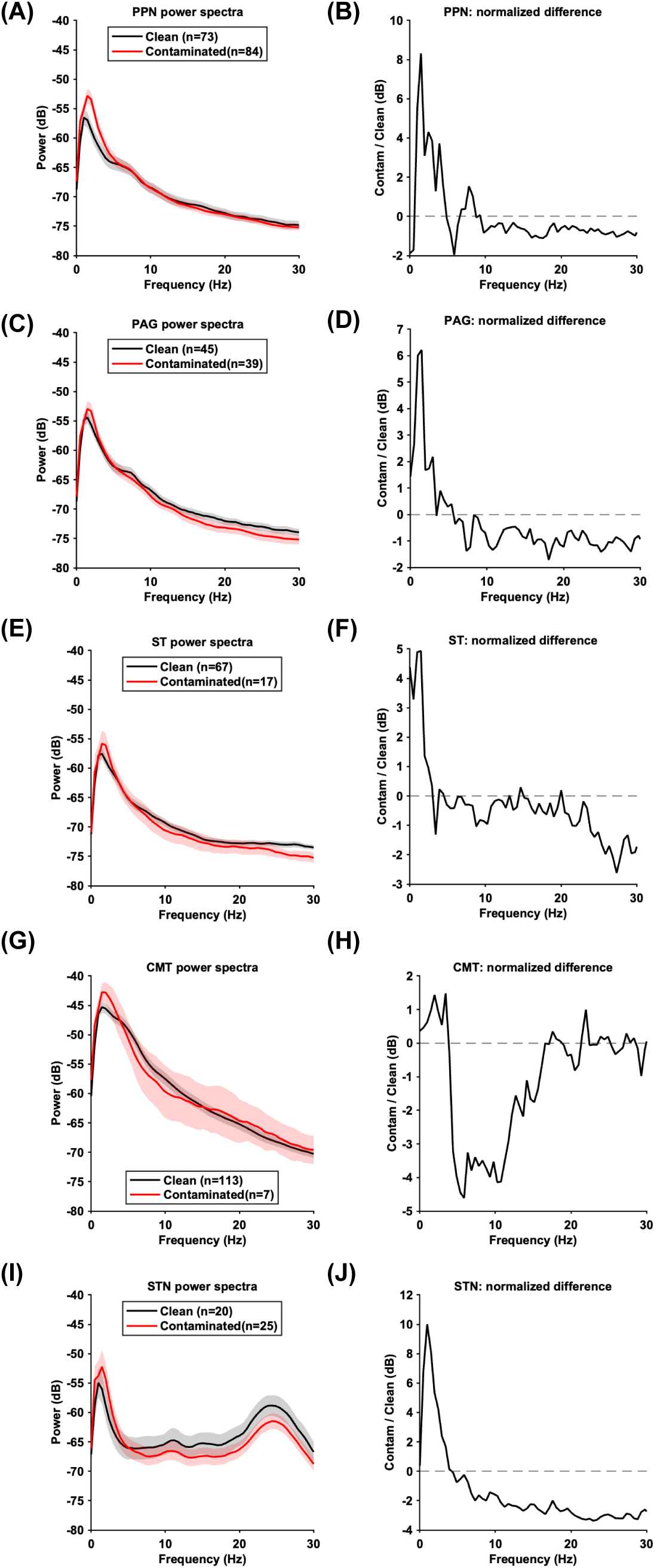
Power spectra of clean and contaminated LFPs. Power spectra and normalized spectral differences in clean and contaminated power spectral density (PSD) for clean and contaminated LFPs for the PPN (A, B), PAG (C,D), ST (E,F), CMT (G,H), STN (I,J). Mean power spectral densities were first calculated by averaging in linear power units. A normalized spectral difference was then computed at each frequency as the logarithmic ratio between contaminated and clean conditions. The PPN, PAG, ST, and STN plots show contamination produced frequency-dependant increases in spectral power with maximal effects in low frequencies. (A-B) This difference was significant for delta frequencies in the PPN (B-H adjusted p = 0.0.021) and (C-D) PAG (B-H adjusted p = 0.013). (E-F) In the ST there were significant differences in beta (B-H adjusted p = 0.011) and gamma (B-H adjusted p = 0.011). (G-H) In the CMT there was significantly different spectral power in the theta (B-H adjusted p = 0.026) and alpha power (B-H adjusted p = 0.026) in contaminated traces. (I-J) There were no significant differences in the STN. B-H adjusted = Benjamini-Hochberg adjusted, PPN = pedunculopontine nucleus, PAG = periaqueductal gray, ST = sensory thalamus, CMT = centromedian nucleus of the thalamus, STN = subthalamic nucleus.

**Table 2.**
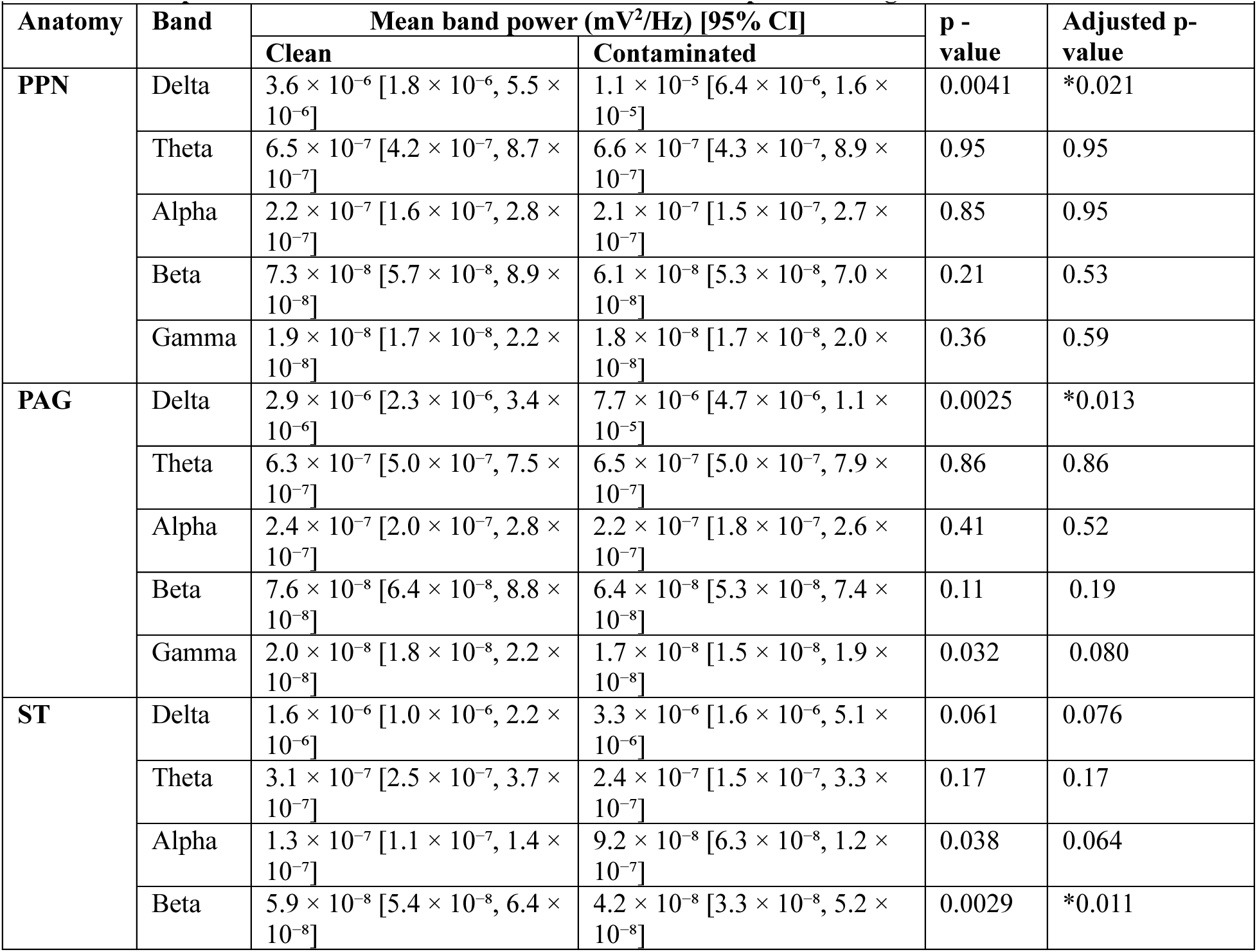
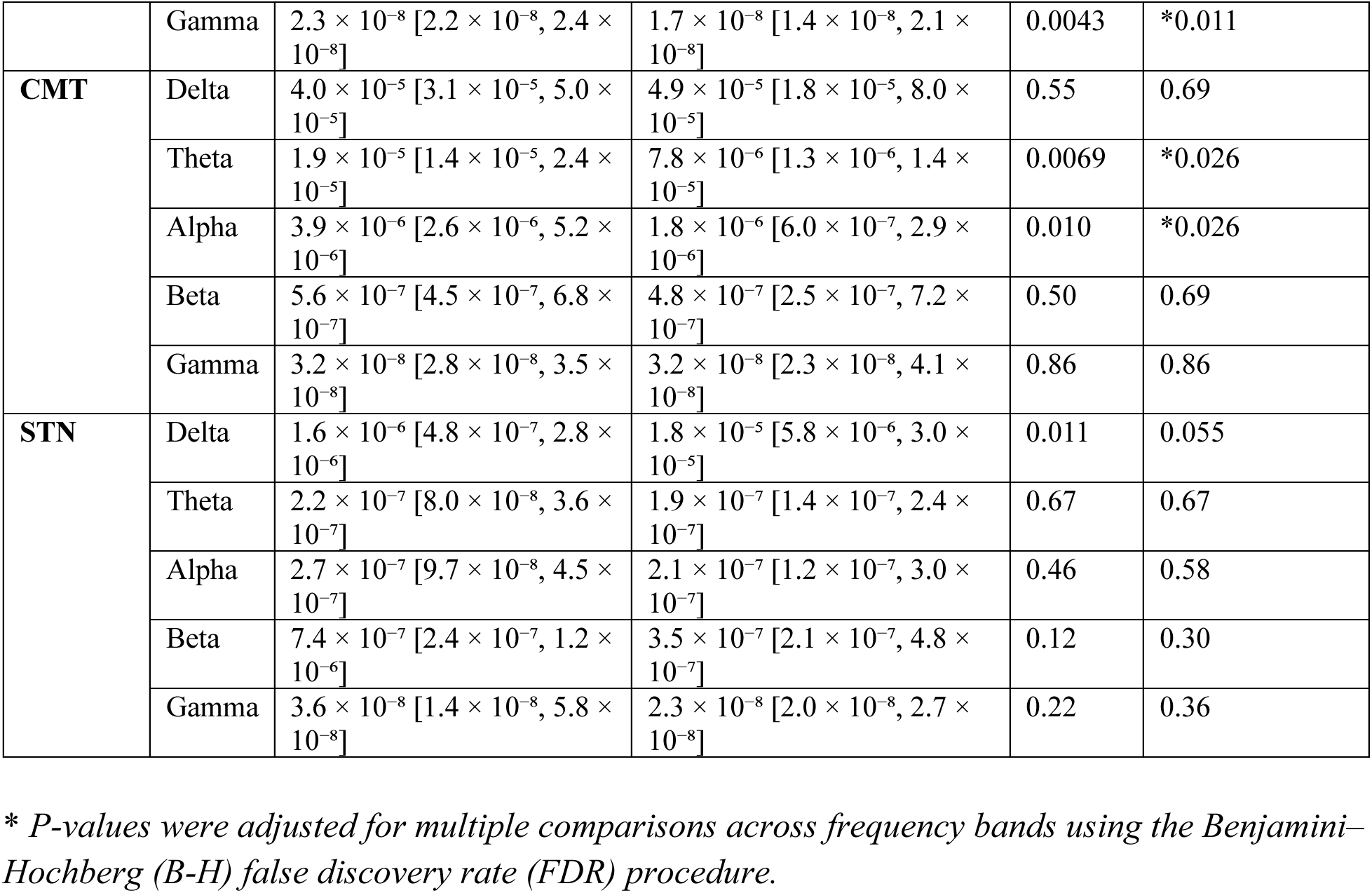
Band power differences in clean vs contaminated LFPs per DBS target.

|  |  |  |  |  |  |
| --- | --- | --- | --- | --- | --- |
| Anatomy | Band | Mean band power (mV<sup>2</sup>/Hz) [95% CI] |  | p - value | Adjusted p-value |
|  |  | Clean | Contaminated |  |  |
| PPN | Delta | $3.6 \times 10^{-6}$ [$1.8 \times 10^{-6}$, $5.5 \times 10^{-6}$] | $1.1 \times 10^{-5}$ [$6.4 \times 10^{-6}$, $1.6 \times 10^{-5}$] | 0.0041 | *0.021 |
| | Theta | $6.5 \times 10^{-7}$ [$4.2 \times 10^{-7}$, $8.7 \times 10^{-7}$] | $6.6 \times 10^{-7}$ [$4.3 \times 10^{-7}$, $8.9 \times 10^{-7}$] | 0.95 | 0.95 |
| | Alpha | $2.2 \times 10^{-7}$ [$1.6 \times 10^{-7}$, $2.8 \times 10^{-7}$] | $2.1 \times 10^{-7}$ [$1.5 \times 10^{-7}$, $2.7 \times 10^{-7}$] | 0.85 | 0.95 |
| | Beta | $7.3 \times 10^{-8}$ [$5.7 \times 10^{-8}$, $8.9 \times 10^{-8}$] | $6.1 \times 10^{-8}$ [$5.3 \times 10^{-8}$, $7.0 \times 10^{-8}$] | 0.21 | 0.53 |
| | Gamma | $1.9 \times 10^{-8}$ [$1.7 \times 10^{-8}$, $2.2 \times 10^{-8}$] | $1.8 \times 10^{-8}$ [$1.7 \times 10^{-8}$, $2.0 \times 10^{-8}$] | 0.36 | 0.59 |
| PAG | Delta | $2.9 \times 10^{-6}$ [$2.3 \times 10^{-6}$, $3.4 \times 10^{-6}$] | $7.7 \times 10^{-6}$ [$4.7 \times 10^{-6}$, $1.1 \times 10^{-5}$] | 0.0025 | *0.013 |
| | Theta | $6.3 \times 10^{-7}$ [$5.0 \times 10^{-7}$, $7.5 \times 10^{-7}$] | $6.5 \times 10^{-7}$ [$5.0 \times 10^{-7}$, $7.9 \times 10^{-7}$] | 0.86 | 0.86 |
| | Alpha | $2.4 \times 10^{-7}$ [$2.0 \times 10^{-7}$, $2.8 \times 10^{-7}$] | $2.2 \times 10^{-7}$ [$1.8 \times 10^{-7}$, $2.6 \times 10^{-7}$] | 0.41 | 0.52 |
| | Beta | $7.6 \times 10^{-8}$ [$6.4 \times 10^{-8}$, $8.8 \times 10^{-8}$] | $6.4 \times 10^{-8}$ [$5.3 \times 10^{-8}$, $7.4 \times 10^{-8}$] | 0.11 | 0.19 |
| | Gamma | $2.0 \times 10^{-8}$ [$1.8 \times 10^{-8}$, $2.2 \times 10^{-8}$] | $1.7 \times 10^{-8}$ [$1.5 \times 10^{-8}$, $1.9 \times 10^{-8}$] | 0.032 | 0.080 |
| ST | Delta | $1.6 \times 10^{-6}$ [$1.0 \times 10^{-6}$, $2.2 \times 10^{-6}$] | $3.3 \times 10^{-6}$ [$1.6 \times 10^{-6}$, $5.1 \times 10^{-6}$] | 0.061 | 0.076 |
| | Theta | $3.1 \times 10^{-7}$ [$2.5 \times 10^{-7}$, $3.7 \times 10^{-7}$] | $2.4 \times 10^{-7}$ [$1.5 \times 10^{-7}$, $3.3 \times 10^{-7}$] | 0.17 | 0.17 |
| | Alpha | $1.3 \times 10^{-7}$ [$1.1 \times 10^{-7}$, $1.4 \times 10^{-7}$] | $9.2 \times 10^{-8}$ [$6.3 \times 10^{-8}$, $1.2 \times 10^{-7}$] | 0.038 | 0.064 |
| | Beta | $5.9 \times 10^{-8}$ [$5.4 \times 10^{-8}$, $6.4 \times 10^{-8}$] | $4.2 \times 10^{-8}$ [$3.3 \times 10^{-8}$, $5.2 \times 10^{-8}$] | 0.0029 | *0.011 |
| | Gamma | $2.3 \times 10^{-8}$ [$2.2 \times 10^{-8}$, $2.4 \times 10^{-8}$] | $1.7 \times 10^{-8}$ [$1.4 \times 10^{-8}$, $2.1 \times 10^{-8}$] | 0.0043 | *0.011 |
| **CMT** | Delta | $4.0 \times 10^{-5}$ [$3.1 \times 10^{-5}$, $5.0 \times 10^{-5}$] | $4.9 \times 10^{-5}$ [$1.8 \times 10^{-5}$, $8.0 \times 10^{-5}$] | 0.55 | 0.69 |
| | Theta | $1.9 \times 10^{-5}$ [$1.4 \times 10^{-5}$, $2.4 \times 10^{-5}$] | $7.8 \times 10^{-6}$ [$1.3 \times 10^{-6}$, $1.4 \times 10^{-5}$] | 0.0069 | *0.026 |
| | Alpha | $3.9 \times 10^{-6}$ [$2.6 \times 10^{-6}$, $5.2 \times 10^{-6}$] | $1.8 \times 10^{-6}$ [$6.0 \times 10^{-7}$, $2.9 \times 10^{-6}$] | 0.010 | *0.026 |
| | Beta | $5.6 \times 10^{-7}$ [$4.5 \times 10^{-7}$, $6.8 \times 10^{-7}$] | $4.8 \times 10^{-7}$ [$2.5 \times 10^{-7}$, $7.2 \times 10^{-7}$] | 0.50 | 0.69 |
| | Gamma | $3.2 \times 10^{-8}$ [$2.8 \times 10^{-8}$, $3.5 \times 10^{-8}$] | $3.2 \times 10^{-8}$ [$2.3 \times 10^{-8}$, $4.1 \times 10^{-8}$] | 0.86 | 0.86 |
| **STN** | Delta | $1.6 \times 10^{-6}$ [$4.8 \times 10^{-7}$, $2.8 \times 10^{-6}$] | $1.8 \times 10^{-5}$ [$5.8 \times 10^{-6}$, $3.0 \times 10^{-5}$] | 0.011 | 0.055 |
| | Theta | $2.2 \times 10^{-7}$ [$8.0 \times 10^{-8}$, $3.6 \times 10^{-7}$] | $1.9 \times 10^{-7}$ [$1.4 \times 10^{-7}$, $2.4 \times 10^{-7}$] | 0.67 | 0.67 |
| | Alpha | $2.7 \times 10^{-7}$ [$9.7 \times 10^{-8}$, $4.5 \times 10^{-7}$] | $2.1 \times 10^{-7}$ [$1.2 \times 10^{-7}$, $3.0 \times 10^{-7}$] | 0.46 | 0.58 |
| | Beta | $7.4 \times 10^{-7}$ [$2.4 \times 10^{-7}$, $1.2 \times 10^{-6}$] | $3.5 \times 10^{-7}$ [$2.1 \times 10^{-7}$, $4.8 \times 10^{-7}$] | 0.12 | 0.30 |
| | Gamma | $3.6 \times 10^{-8}$ [$1.4 \times 10^{-8}$, $5.8 \times 10^{-8}$] | $2.3 \times 10^{-8}$ [$2.0 \times 10^{-8}$, $2.7 \times 10^{-8}$] | 0.22 | 0.36 |

\* *P-values were adjusted for multiple comparisons across frequency bands using the Benjamini–Hochberg (B-H) false discovery rate (FDR) procedure.*

### Artifact characteristics

To examine the characteristics of ‘pure’ pulsatile signal independent of background activity, we attempted to extract an artifact template using the PulsAr algorithm (see Methods). A sample output of the first and second cross correlations and pulsatile signal template extraction is shown in figure 6A-B. For each contaminated LFP, the pulsatile template was extracted, frequency spectra inspected, and templates stitched together in repeats to form a pulsatile template string independent of underlying LFP activity (figure 6 C, E, G, I, K). Spectral analysis of the templates (figure 6 D,F,H,J,L) showed the greatest power in lower (delta) frequencies, with spectral power tailing off towards higher frequencies and flattening out around 10-15Hz. In the STN only, a peak in the beta frequency range was observed, indicating a potential influence of STN beta activity on the template extraction.

**Figure 6.**
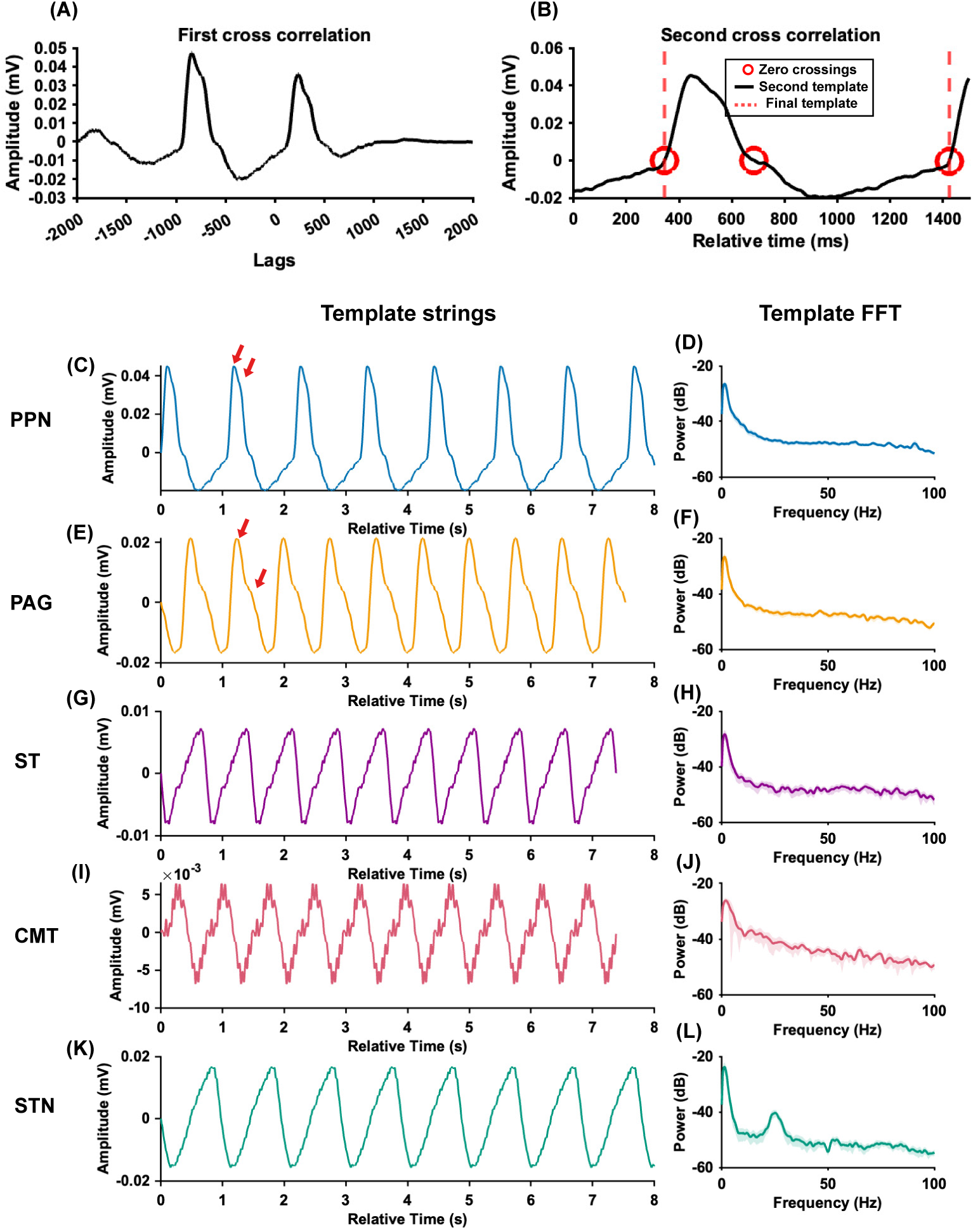
Pulsatile signal detection, extraction, and frequency content evaluation. (A) A sample output from the extraction element of the PulsAr algorithm. In the first cross correlation, the signal is cross correlated with itself to identify areas of high correlation, a repeating shape in the signal. The area with highest correlation is mapped back to the signal to extract a first template shape. (B) The first template is cross correlated with the original signal again and areas of highest correlation are extracted and averaged together to form the second template. For later processing, and to prevent square waves, we select where the signal crosses zero. The final template is then used to make strings of artifact. (C) Pulsatile waveform strings extracted from a sample PPN LFP. (D) Mean +/- 95% CI of the fast Fourier transform (FFT) of templates extracted from all contaminated PPN signals. Red arrows indicate primary and secondary peak morphology resembling P1 and P2 in an intracranial pressure waveform. (E) Pulsatile waveform strings extracted from a sample PAG LFP. Red arrows indicate primary and secondary peak morphology resembling P1 and P2 in an intracranial pressure waveform. (F) Mean +/- 95% CI of the FFT of templates extracted from all contaminated PAG signals. (G) Pulsatile waveform strings extracted from a sample ST LFP. (H) Mean +/- 95% CI of the FFT of templates extracted from all contaminated ST signals. (I) Pulsatile waveform strings extracted from a sample CMT LFP. (J) Mean +/- 95% CI of the FFT of templates extracted from all contaminated CMT signals. (K) Pulsatile waveform strings extracted from a sample STN LFP. (L) Mean +/- 95% CI of the FFT of templates extracted from all contaminated STN signals. PPN = pedunculopontine nucleus, PAG = periaqueductal grey, ST = sensory thalamus, CMT = centromedian nucleus of the thalamus, STN = subthalamic nucleus.

We sought to determine why PulsAr detected contamination in CMT LFPs despite no visual pulsatile waveform. We compared RMS amplitudes of clean, contaminated, and extracted pulsatile signal templates from LFPs within each brain region (figure 7). Mann-Whitney U tests between clean and contaminated signals showed significant differences in the PPN (p = 2.5x10^-7^) and PAG (p = 0.0018) with RMS amplitude in the contaminated signals significantly higher than clean signals (figure 7A). There were no significant differences in the ST, CMT, or STN (p > 0.05). There were no significant differences when comparing RMS amplitude of pulsatile templates between DBS targets (Kruskal-Wallis, p = 0.16), figure 7B. There were also no significant differences in in the proportion of total contaminated LFP signal attributable to pulsatile signal energy as assessed by comparing contamination indices between DBS targets (Kruskal-Wallis, p = 0.11, figure 7C). Nevertheless, RMS amplitude of clean CMT LFPs appears to trend high compared to the other DBS targets (figure 7A), while the RMS amplitude of the extracted pulsatile template (figure 7B) is not larger than others. For the CMT, this suggests that a pulsatile component, if present, is small compared to endogenous signal potentially making it visually undetectable. This is also reflected in a lower but non-significant contamination index value in the CMT.

**Figure 7.**
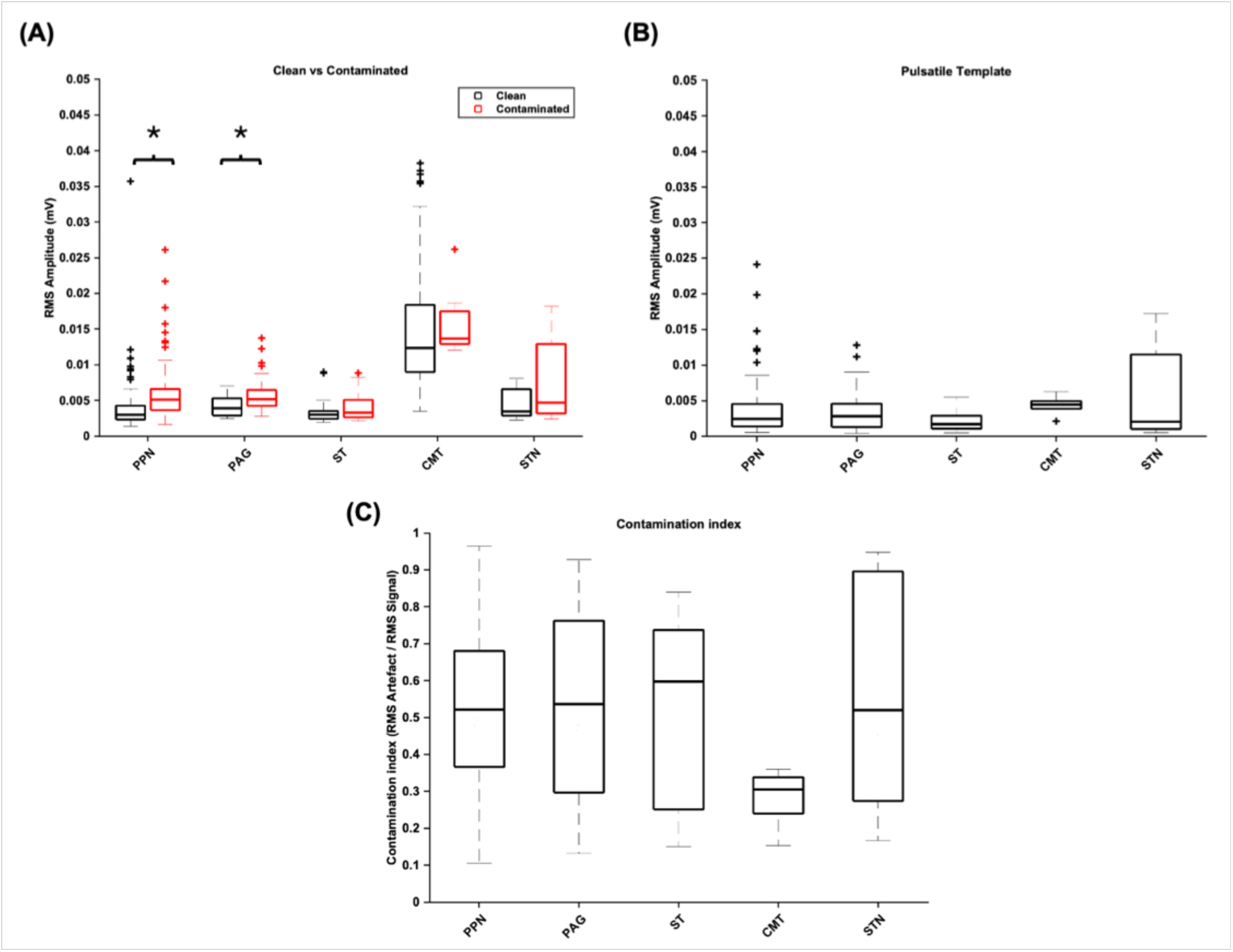
Comparative RMS amplitudes of signals and extracted pulsatile templates. Boxplots show the median (center line), 25th–75th percentiles (box), and minimum/maximum values within 1.5× inter-quartile range (whiskers). Outliers are shown as individual points. (A) RMS amplitudes of clean and contaminated LFPs from each DBS target region. Mann-Whitney U tests between clean and contaminated signals showed significant differences in the PPN (p = 2.5x10^-7^) and PAG (p = 0.0018) with RMS amplitude in the contaminated signals significantly higher than clean signals. There were no significant differences in the ST, CMT, or STN. (B) RMS amplitudes of isolated pulsatile templates from each DBS target region. There were no significant differences between DBS target regions (Kruskal-Wallis, p = 0.16). (C) Contamination index showing signal distortion by pulsatile signal for each DBS target region. There were no significant differences between each DBS target region (Kruskal-Wallis, p = 0.11). *Indicates significance.

To investigate the extent to which filtering might mask or mitigate the pulsatile signal, we applied high pass filters with a range of cut offs to an extracted pulsatile template string as well as a raw contaminated PPN LFP. Figure 8A shows that increasing high pass filter cut off progressively decreases the large amplitude pulsatile component of the template string, but that higher frequency components remain even at higher cutoffs. When applied to a source original contaminated PPN LFP, the remaining spectral components of the pulsatile signal become difficult to visually detect amongst background oscillatory neural activity (figure 8B). To investigate the feasibility of using a linear subtraction approach to mitigate the effect of the pulsatile signal on LFP biomarker extraction we also characterised the stability of the signal over time. In two 40-second PPN recordings, sliding-window analysis showed how the RMS amplitude of the pulsatile component can fluctuate even over relatively short timescales (figure 8C), indicating that the pulsatile signal would affect LFP biomarker estimates differently at different time points.

**Figure 8.**
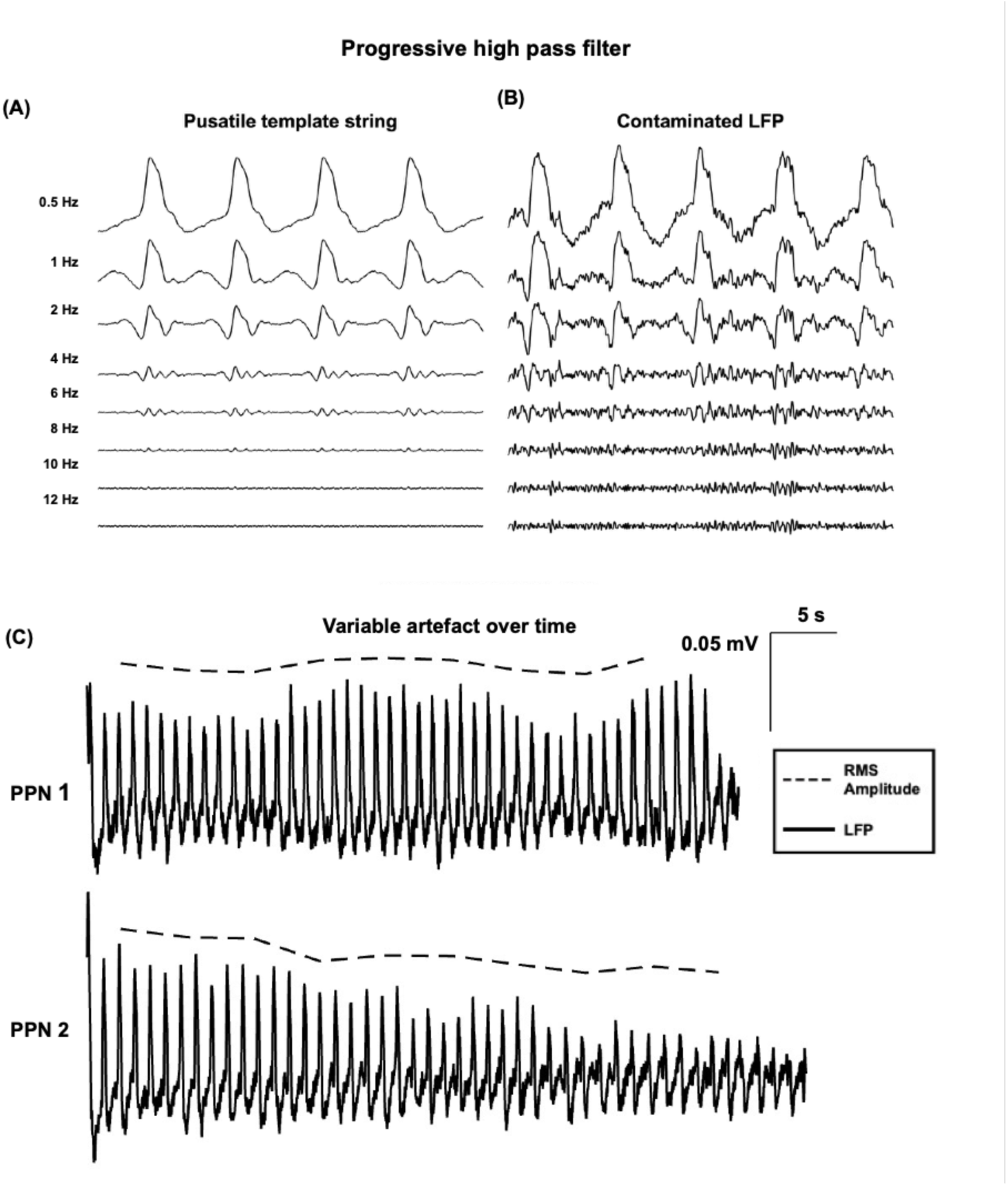
Challenges for pulsatile signal mitigation. (A) Application of sequentially larger high pass filter cut offs to the pulsatile template string with templates extracted from a contaminated PPN LFP. The amplitude of the template peak decreases with higher cut-offs. (B) The original contaminated LFP with the same high pass filter cut-offs show that as the pulsatile signal amplitude decreases it becomes visually challenging to detect against the background endogenous activity. (C) RMS amplitudes of pulsatile templates extracted are not necessarily stable over time. Two sample contaminated signals are presented. RMS amplitude was calculated for five second segments using a sliding window on the contaminated signal. The RMS amplitude follows the observed pulsatile signal within the LFP.

**Table 3.** RMS amplitude comparisons between extracted pulsatile signal templates, and clean and contaminated LFPs.

| DBS Target | Classification | Mean RMS (mV) +/- 95% CI | Clean vs contaminated (Mann-Whitney U) | Pulsatile Template (Kruskal-Wallis) | Contamination Index (Kruskal-Wallis) |
| --- | --- | --- | --- | --- | --- |
| PPN | Clean | 0.0041 (0.0031–0.0052) | *2.55x10<sup>-7</sup> | H(4) = 6.5, p = 0.16 | H (4) = 7.6, p = 0.11 |
|  | Contaminated | 0.0061 (0.0052–0.0070) |  |  |  |
|  | Pulsatile Template | 0.0037 (0.0028–0.0046) |  |  |  |
|  | Contamination Index | 0.53 |  |  |  |
| PAG | Clean | 0.0042 (0.0038–0.0045) | *0.0018 |  |  |
|  | Contaminated | 0.0058 (0.0049–0.0066) |  |  |  |
|  | Pulsatile Template | 0.0035 (0.0025–0.0046) |  |  |  |
|  | Contamination Index | 0.54 |  |  |  |
| ST | Clean | 0.0032 (0.0029–0.0035) | 0.17 |  |  |
|  | Contaminated | 0.0040 (0.0031–0.0050) |  |  |  |
|  | Pulsatile Template | 0.0022 (0.0014–0.0029) |  |  |  |
|  | Contamination Index | 0.52 |  |  |  |
| CMT | Clean | 0.015 (0.013–0.016) | 0.25 |  |  |
|  | Contaminated | 0.016 (0.011–0.020) |  |  |  |
|  | Pulsatile Template | 0.0043 (0.0031–0.0055) |  |  |  |
|  | Contamination Index | 0.28 |  |  |  |
| STN | Clean | 0.0045 (0.0030, 0.0060) | 0.14 |  |  |
|  | Contaminated | 0.0077 (0.0049, 0.010) |  |  |  |
|  | Pulsatile Template | 0.0057 (0.0027, 0.0087) |  |  |  |
|  | Contamination Index | 0.55 |  |  |  |

*\*Indicates statistical significance*

## DISCUSSION

Here we described and characterized a pulsatile signal present in intracranial LFP recordings from DBS leads. We showed that this signal affects recordings across multiple DBS targets and can particularly affect estimates of lower frequency LFP power. Our detection and template reconstruction algorithms enable screening and reconstruction of this artifact without access to ground truth ECG data, providing practical tools for assessing data quality. In the context of CL-DBS, the pulsatile cardiac signal may interfere with neural biomarkers of interest, disrupting adaptive control. Screening DBS recordings for pulsatile contamination is therefore essential for both data interpretation and safe CL-DBS implementation.

### Pulsatile signal prevalence

Our work expands on prior observations of cardiac pulse-synchronous contamination in neural recordings [19,22]. We identified pulsatile signals across multiple DBS targets, neurological conditions, and in two recording setups (a cranially mounted IPG and an externalized lead recording system). Furthermore, we developed an ECG-independent classifier, PulsAr, that reliably identifies contaminated LFPs and extracts representative pulsatile waveform templates. PulsAr showed high sensitivity and specificity relative to visual ratings, which are stable to class imbalance. Frequency-dependent metrics such as accuracy and precision were influenced by the unequal distribution of clean and contaminated traces.

Across montage sweeps, 33.0% of LFPs were contaminated, increasing to 43.1% when excluding the largely clean CMT. In other areas, contamination patterns varied across contacts and sessions. This instability complicates mitigation strategies for CL-DBS based on constant correction factors or contact selection. Because contamination occurred across diseases, age groups, and anatomic areas, the pulsatile signal likely reflects a brain-wide phenomenon across populations.

Although PulsAr identified contamination in a small number of CMT traces, no clear pulsatile waveform was visible. RMS amplitude analysis suggests this may reflect masking by relatively large endogenous CMT signals. Another possibility is PulsAr detection of other repetitive patterns, such as spike-and-wave discharges [26]. Simultaneous LFP–ECG recordings will be necessary to determine whether the CMT contamination is cardiac-related. Differences in surgical technique, such as the use of a guide tube that remained in place in the CADET Pilot trial (Renishaw neuroguide^TM^ DBS electrode delivery system, UK) may also reduce movement of the electrode and therefore LFP contamination.

We considered why this pulsatile signal may have gone previously unrecognized. High pass filters on commercial systems like Medtronic’s Percept^TM^ PC device and Neuropace’s RNS system may remove the more visually obvious components of the pulsatile artifact [23–25]. Our 0.5 Hz high-pass drift removal filter preserved lower-frequency content and enabled clear visualization and detection of the pulsatile waveform. The PulsAr detector was intentionally designed to identify signals repeating at heart-rate frequencies (0.83–2 Hz).

### Pulsatile signal characteristics

The absence of an ECG QRS signature aligns with prior modeling predictions that cranially mounted systems should not see interference from the cardiac dipole [17]. Instead, the pulsatile signal showed a morphology similar to intracranial pressure (ICP) waveforms, with a primary and sometimes a secondary peak [27]. The extracted pulsatile signal contained frequencies up to at least 5-10 Hz and was variable over time, complicating removal without data loss. A peak in the beta band was seen in the extracted pulsatile template from the STN, which might be due to prominence of beta in this region interacting with our template extraction algorithm leading to unintentional phase-alignment, as we would not expect an ICP-driven waveform to have a strong beta component.

Consistent with the dominant low frequency content of the heart pulse signal, PPN and PAG had significantly different delta band power between clean and contaminated signals. Delta band activity is implicated in various physiologic and pathologic brain states [28]. The presence of pulsatile contamination must be considered in future LFP studies targeting these regions. Contaminated CMT recordings showed a small decrease in theta and alpha band power, suggesting a non-additive interaction of artifact and true signal. This could reflect increased likelihood of artifact detection against certain types of background activity; however, interpretation is limited by the small artifact magnitude with respect to the signal and small number of contaminated traces.

### Mechanism of pulsatile artifact generation

We hypothesize that cardiac-driven brain pulsatility generates pulse-synchronous LFP components via electromechanical coupling (figure 9) [29]. Brain pulsatility arises from cyclic changes in vascular and cerebrospinal fluid volumes that compensate for each other to maintain stable intracranial pressure. Brain pulsations coupled to the cardiac cycle can be seen in the fontanelle of newborns, on phase-gated MRI, and intraoperative visualization [27,30–33]. The morphologic similarity of the waveform in contaminated tracts (figure 3A&C) to an ICP waveform, with a first (P1) and sometimes second peak (P2), supports a mechanical origin [27]. The fact that an analogous P1 and P2 were captured in the extracted pulsatile templates suggest it is a common feature captured during the averaging process (6C&E). Higher contamination rates in the brainstem-adjacent regions, PPN and PAG, is consistent with known spatial gradients in brain pulsatility [34].

**Figure 9.**
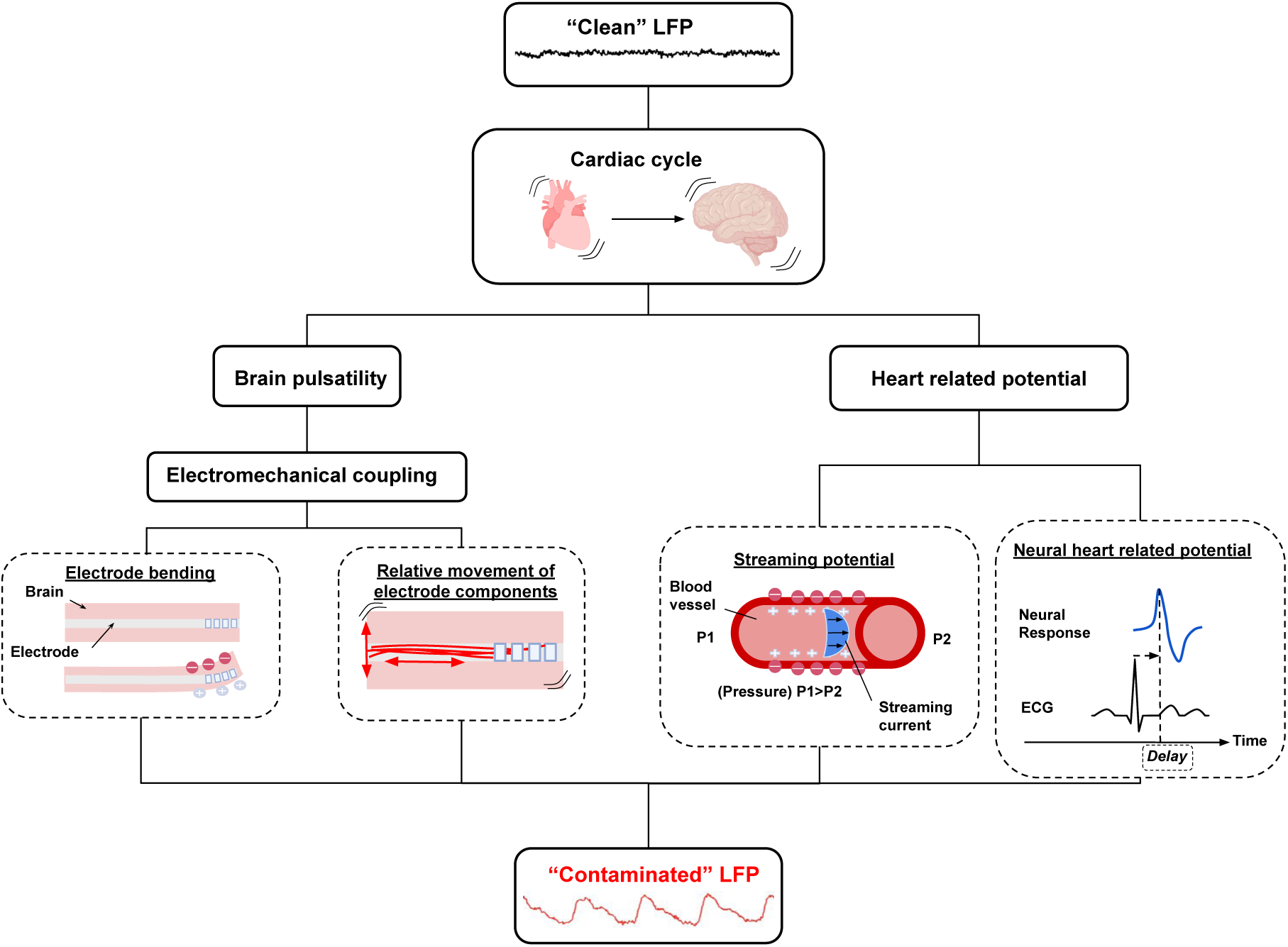
Mechanisms of pulsatile signal generation related to the cardiac cycle. A schematic showing the hypothesized process of generating an LFP contaminated with pulsatile signal. The brain pulses during the cardiac cycle due to systolic arterial blood volume causing compensatory changes in intracranial CSF volume. We hypothesize that brain pulsatility through electromechanical coupling causes the generation of the pulsatile signal. This could be due to electrode bending and impedance changes, the relative movement of electrode components (such as the wires within the lead causing a relative change in the perceived voltage). Alternative hypotheses include a physiology heart beat related potential like a streaming potential, or heart beat evoked neural activity.

An additional mechanical mechanism may include electrode component micromotion. Electromechanical coupling differences due to anatomic location or proximity to vessels or CSF spaces, may contribute and require further investigation. Given the ‘macro’ scale of brain pulsations, it is interesting that pulsatile artifact is not rejected as a common mode and is present in bipolar recordings. Differential amplification may convert small asymmetries in electrode deformation into measurable voltage differences, explaining variable contamination patterns.

While electromechanical interactions between the pulsating tissue and the recording leads might account for the pulsatile signal, it is also possible that the ‘true’ LFP in fact does fluctuate according to the cardiac cycle, via neural activity synchronised to the cardiac cycle or through streaming potentials from pulsatile ionic blood flow (figure 9)[35,36]. For instance, heart cycle related effects such as heartbeat-evoked potentials (HEPs) have described in other neural recordings such as electroencephalograms (EEG) and electrocorticograms (ECoG), though these often require triggered averaging to be observable and differ in shape from the ICP-like waveform observed here[35,37–39]. At the neuronal level, it has been shown that mechanosensitive ion channels can mediate responses to cardiovascular pressure pulsations and that these can entrain local neuronal activity [40]. There is less literature on streaming potential effects in the brain but as these would be directly driven by blood flow fluctuations these would more closely mirror the ICP waveform. As these mechanisms are not mutually exclusive and could all be contributing factors, it may prove difficult to disentangle their individual contributions *in vivo*.

### Clinical implications and potential use as a tool

Pulsatile contamination poses challenges for CL-DBS. Like intracranial pulsatility [13,27], the pulsatile signal exhibits oscillations beyond the fundamental heart-rate frequency. We show high-pass filtering alone may be insufficient to remove pulsatile contamination given that in our spectrograms the pulsatile signal extends to >10 Hz and small rhythmic components remained visible with higher filter cutoffs when used on pure artifact strings (figures 3&8). This signal may affect LFP analyses used for clinical decisions, and in a closed-loop system could lead to false threshold crossings and unintended stimulation. Future studies of low frequency oscillatory activity using DBS systems should be aware that LFPs may be affected by heart rate or intracranial pressure and compliance (also affected by e.g. posture). This is particularly important in long term band power tracking where the underlying full spectrum timeseries is not saved, preventing post-hoc screening for cardiac contamination. PulsAr provides an objective, ECG independent screening tool and could support online monitoring or contact selection in the future.

However, mitigating or removing the ‘artifact’ may not be appropriate if this pulsatile signal is in fact a true feature of the local field potential. In that case, the question is to what extent this signal interacts with ongoing neural activity and whether it should be removed in analysis to improve estimates of superimposed oscillatory activity biomarkers. Future work should therefore determine if this signal is neural in nature, and if so, activity whether it shows non-linear interactions with other neural oscillations of interest. An additional intriguing possibility is that the PulsAr detector and pulsatile signals in LFPs could be repurposed as a clinical tool, for example as a heart-rate estimator for sleep staging, seizure-related tachycardia detection, or as a proxy for intracranial pressure dynamics.

### Limitations and future directions

PulsAr is designed to detect large amplitude, regularly repeating signals at physiological heart rate frequencies, and therefore by definition is vulnerable to false positives if such signals are present in underlying neural activity. Simultaneous ECG-LFP recordings are needed to validate PulsAr as an intracranial heart rate detector for specific clinical targets and to control for this effect. In our case, simultaneous ECG and LFP recordings will be required to confirm whether CMT contamination detected by PulsAr is cardiac related. Further longitudinal data is required to determine if there is chronic contamination patterns among contact pairs. ICP and brain pulsatility is known to change with position and during sleep and it would be interesting to investigate the effect of position on the pulsatile signal [31,33,41–43]. Finally, determining the electromechanical mechanism generating the pulsatile signal might allow for design of DBS technology that is less sensitive to cardiac pulse contamination.

## CONCLUSIONS

We demonstrate that heart pulse signal is present in recordings from DBS leads across multiple targets and clinical populations. While the most obvious components may be filtered out by commercial device streaming filters, it affects frequencies above heart rate and could affect biomarkers used for clinical decisions. Our ECG-independent detector reliably identifies and extracts this signal. Broadband spectral content and temporal variability complicate automated removal. Recognising and mitigating pulsatile contamination is essential for accurate sensing and safe closed-loop therapy. Further work is needed to clarify its physiological origin and clinical significance.

## ACKNOWLEDGMENTS

We acknowledge contributions of the clinical teams involved in trial design and data collection. For the MINDS trial, we thank Alceste Deli, Robert Toth, Mayela Zamora, and Amir P. Divanbeighi Zand. For EPIONE, we acknowledge Victoria Marks, Rachel Crockett, John Eraifej, and Martin Gillies. For CADET Pilot, we thank Marios Kaliakatsos, Antonio Valentin, Victoria Marks, John Fleming, Sarah Carter, and Friederike Moeller.

## AUTHOR CONTRIBUTIONS

Conceptualization: KT, JvR, TD; Methodology: KT and JvR; Data Collection: TD, MT, AG, JN, RP, JvR; Data Analysis: KT, JvR; Writing: KT, JvR; Review & Editing: KT, JvR, TD, MT, AG, WJN, RP

## DATA AVAILABILITY

Code will be released on manuscript acceptance following peer review. The dataset is not yet openly available as it is being used in ongoing projects. Due to data protection regulations access to the raw data would need to be in a trusted research environment as part of a collaboration. Please contact.

## USE OF GENERATIVE AI

ChatGPT 5.2 was used to edit code, for manuscript language editing, and improvement of manuscript clarity.

